# Oligodendrocyte precursor cells prune axons in the mouse neocortex

**DOI:** 10.1101/2021.05.29.446047

**Authors:** JoAnn Buchanan, Leila Elabbady, Forrest Collman, Nikolas L. Jorstad, Trygve E. Bakken, Carolyn Ott, Jenna Glatzer, Adam A. Bleckert, Agnes L. Bodor, Derrick Brittan, Daniel J. Bumbarger, Gayathri Mahalingam, Sharmishtaa Seshamani, Casey Schneider-Mizell, Marc M. Takeno, Russel Torres, Wenjing Yin, Rebecca D. Hodge, Manuel Castro, Sven Dorkenwald, Dodam Ih, Chris S. Jordan, Nico Kemnitz, Kisuk Lee, Ran Lu, Thomas Macrina, Shang Mu, Sergiy Popovych, William M. Silversmith, Ignacio Tartavull, Nicholas L. Turner, Alyssa M. Wilson, William Wong, Jingpeng Wu, Aleksandar Zlateski, Jonathan Zung, Jennifer Lippincott-Schwartz, Ed S. Lein, H. Sebastian Seung, Dwight E. Bergles, R. Clay Reid, Nuno Maçarico da Costa

## Abstract

Neurons in the developing brain undergo extensive structural refinement as nascent circuits adopt their mature form^1^. This transformation is facilitated by the engulfment and degradation of excess axonal branches and inappropriate synapses by surrounding glial cells, including microglia and astrocytes^2,3^. However, the small size of phagocytic organelles and the complex, highly ramified morphology of glia has made it difficult to determine the contribution of these and other glial cell types to this process. Here, we used large scale, serial electron microscopy (ssEM) with computational volume segmentation to reconstruct the complete 3D morphologies of distinct glial types in the mouse visual cortex. Unexpectedly, we discovered that the fine processes of oligodendrocyte precursor cells (OPCs), a population of abundant, highly dynamic glial progenitors^4^, frequently surrounded terminal axon branches and included numerous phagolysosomes (PLs) containing fragments of axons and presynaptic terminals. Single- nucleus RNA sequencing indicated that cortical OPCs express key phagocytic genes, as well as neuronal transcripts, consistent with active axonal engulfment. PLs were ten times more abundant in OPCs than in microglia in P36 mice, and declined with age and lineage progression, suggesting that OPCs contribute very substantially to refinement of neuronal circuits during later phases of cortical development.

## MAIN

OPCs emerge from several germinal zones in late prenatal development following the sequential generation of neurons and astrocytes, migrate into the expanding cortex and then proliferate to establish a grid-like distribution, with individual cells occupying distinct territories. Genetic fate tracing and time lapse imaging *in vivo* have demonstrated that these progenitors play a critical role in generating oligodendrocytes and thus myelin throughout the central nervous system (CNS)^5^. However, OPCs are present in some cortical regions weeks before oligodendrogenesis begins and extend highly ramified processes that developing neurons. These progenitors express a diverse set of neurotransmitter receptors and form direct, functional synapses with excitatory and inhibitory neurons.^6,7^ These features have traditionally been viewed through the perspective of oligodendrogenesis^8^, as other functions for these ubiquitous glial cells have not been clearly established. Our knowledge about the structure and function of OPCs has been severely limited by incomplete ultrastructural information, because their fine, highly branched processes are difficult to unambiguously identify in EM studies without complete reconstructions to connect them to identifiable somata. Modern computational volume EM methods offer new opportunities to increase our understanding of brain ultrastructure^9–12^, particularly for morphologically complex, highly dynamic glial cells like OPCs. Here, we used two densely segmented and reconstructed datasets of mouse visual cortex, ages P36 (Fig. 1a) and P49, to perform a detailed morphometric analysis and quantification of the anatomical features of somata, processes, and organelles of OPCs at this highly dynamic phase of neocortical maturation to help define their roles in the developing brain.

**Figure 1:**
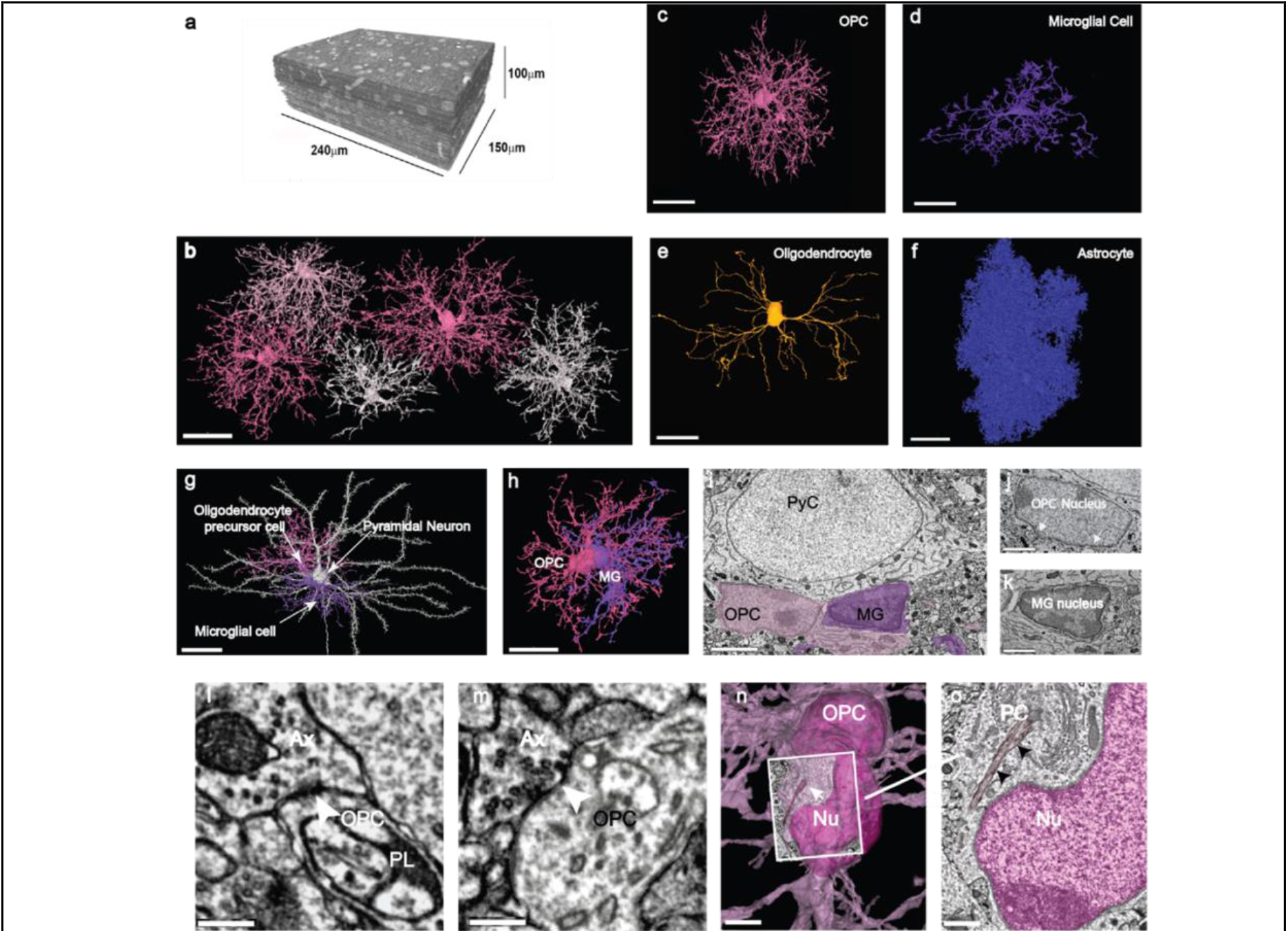
Distinct structural features of OPCs in the developing visual cortex. **a,** ssEM reconstruction of 100 μm^3^ volume of layer 2/3 mouse visual cortex (P36). **b,** 3D reconstructions of a subset of OPCs in P36 dataset, showing discrete territories and tiling. Scale bar, 30 μm. **c**, 3D rendering of an OPC from P36 dataset showing extensive ramifications emanating from the cell soma. **d**, 3D rendering of a microglial cell shows its shorter, less branched processes and elongated and flattened soma. **e**, 3D rendering of a mature myelinating oligodendrocyte in P36 dataset has a smooth and ovoid shaped soma. Note, only the soma and cytoplasmic processes without myelin sheaths are shown. **f**, 3D rendering of an astrocyte in P36 dataset showing its densely packed cytoplasmic protrusions. Scale bars c-f, 20 μm. **g,** 3D rendered pyramidal neuron (white) with an OPC (pink) and a microglial cell (MG, purple) both in satellite positions. Scale bar, 30 μm. (See also Extended Data Video 1) **h,** 3D rendering of the same two glial cells in **g** shows their close association and intermingling of branches. Scale bar, 20 μm. **i,** Ultrathin section slice through an OPC soma (pink) and microglial cell soma (purple). Scale bar, 3 μm. **j,** Ultrathin section slice of an OPC nucleus with dense rim of heterochromatin and ruffled edge (arrows). Scale bar,1.5 μm. **k,** Ultrathin section slice of a microglial nucleus showing its dense heterochromatin throughout. Scale bar, 1.5 μm. **l-m,** Axons (Ax) making synaptic contacts with OPC processes (see also Extended Data Fig.1k). Scale bar, 300 nm. **n,** The soma of an OPC_(dark pink) bears a primary cilium_(arrow) adjacent to the nucleus_(Nu). Scale bar, 3 μm. **o**, Ultrathin slice of the boxed area in **n** showing the primary cilium (PC)(arrows) close to the OPC nucleus (Nu, dark pink). Scale bar, 750 nm. (See also Extended Data Video 2, for details on how to explore public data).

### Structural features of OPCs revealed by large scale serial EM

Past EM ultrastructural studies defined several common morphological characteristics of OPCs, such as their bean-shaped nucleus containing low heterochromatin and presence of centrioles in their cytoplasm^13^, consistent with their proliferative progenitor state.^14^. Originally referred to as a multipotential type of glia^4^, and often described as NG2 cells^15,16^ in reference to their expression of the proteoglycan NG2, early EM investigations often focused on their cellular responses to injury and the close association of their processes with synapses and degenerating nerve fibres^15,17^. More recent studies indicate that OPC processes form direct synapses with axons and that OPCs contact nodes of Ranvier^6,18^ in naïve animals; however, quantitative analysis of these structural features has been difficult due to the limited sampling ^6,15,17,18^.

The two EM volumes <u>P36 (Fig. 1.a)</u> and P49 we examined in serial sections contained four distinct classes of glia: OPCs (Fig. 1b, c), microglia (Fig. 1d), oligodendrocytes (Fig.1e), and astrocytes (Fig. 1f). OPCs exhibited a ramified form with 15 to 17 highly branched processes extending up to 50 μm radially from the soma (Fig. 1b, c, g, h, Extended Data Fig.1 c-j) that contained numerous filopodia on their tips. 3D renderings of multiple cells revealed that OPCs, similar to astrocytes and microglia ^19,20^, formed a grid-like organization with little overlap between territories of adjacent cells of the same class (Fig. 1b), and also contacted synapses^21,22^. The OPC somata were frequently in a satellite position, like those of microglia and oligodendrocytes^23,24^ (Fig. 1g, Extended Data Fig. 1c-k;Video 1), and were remarkably variable in both size and shape, ranging from elongated or bean shaped to smooth or irregular with a rough surface (Fig. 1b; Extended Data Fig. 1c-j). OPCs were readily distinguished from other glial cell types by these features as well as having: 1) larger nuclei that were elongated and contained less heterochromatin than those of microglia (Fig. 1i-k); 2) processes that were longer and smoother compared to microglia or astrocytes (Fig. 1c, d, f); 3) axons that formed synaptic contacts with their processes (Fig. 1l, m); 4) cytoplasm that was more electron lucent than that of microglia and devoid of glycogen granules that are numerous in astrocytes (Fig. 1m); and 5) the presence of primary cilia, which were not found in microglia, pre-myelinating or mature oligodendrocytes (Fig. 1n,o; Extended Data Fig. 1c-j,1k).

### OPC processes contain numerous lysosomes and phagolysosomes

Previous *in vivo* imaging experiments revealed the dynamic nature of OPC processes, which exhibit continuous branch remodeling as they migrate through the gray matter, making transient interactions with various cellular constituents ^25^. Based on profiles in EM images and associated cellular reconstructions, we find that OPC processes were often found in contact with axons (Fig.2a-d). Unexpectedly, the cytoplasm of these processes often contained numerous membrane-bound organelles that included phagosomes and primary and secondary lysosomes, termed phagolysosomes(PLs)(Fig. 2b,c,e), suggesting that OPCs at this age are engaging in phagocytosis. The process of phagocytosis begins with recognition of a target, followed by phagosome formation (engulfment) and maturation, and then fusion of the phagosome with a lysosome, forming a PL^26^. In both EM volumes, phagosomes were identified as completely internalized, membrane delimited structures. Lysosomes appeared electron dense and measured ~ 500 nm in diameter (Fig. 2b), while PLs were multichambered, large organelles measuring ~ 750 nm (Fig. 2e-n, Extended Data Video 2). Seldom visualized in large numbers by EM, PLs represents the final step of phagocytosis, and constitute a highly acidic compartment capable of destroying ingested elements^26,27^. Mapping the distribution of PLs in one OPC (Fig. 2o) demonstrated that they were widely distributed within the cell, but particularly prevalent near the tips of OPC processes. This finding led us to manually quantify PLs in eight more 3D reconstructed OPCs in P36 dataset for comparison.

**Figure 2:**
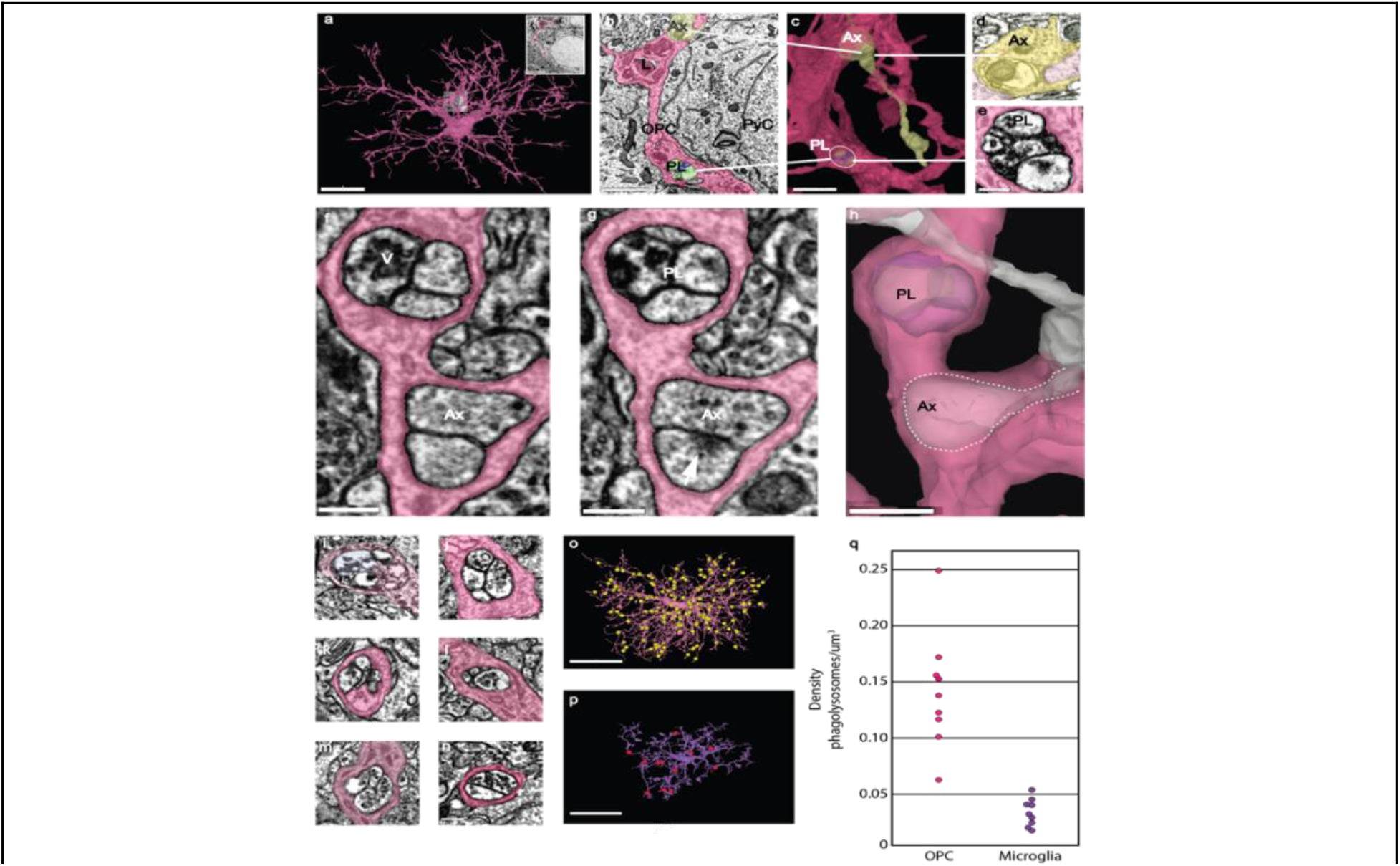
Phagolysosomes are abundant in OPC processes. **a,** 3D rendering of an OPC from the P36 dataset showing ramified branches. Scale bar, 15 μm. Insert shows the same area as in **b**. **b**, Ultrathin section slice of OPC branch containing a phagolysosome (PL), a lysosome (L) and an axon (Ax) passing through the OPC cytoplasm. The branch contacts the adjacent pyramidal cell (PyC). Scale bar, 1.5 μm. **c**, 3D view of the same OPC branch showing the axon (Ax) (yellow) partially encased within the OPC cytoplasm. The same phagolysosome (PL) in **2b** is visible (white circle). Scale bar, 1.5 μm. **d,** Higher magnification view of axon in **c** (yellow). Scale bar, 300 nm**. e,** Higher magnification of phagolysosome (PL) (white line) in **c** highlighting multiple chambers containing cellular debris. Scale bar, 300 nm. **f,** OPC branch from P36 dataset contains a phagolysosome with vesicles (v) and an encapsulated axon (Ax). Scale bar, 300 nm. **g,** A portion of the same branch showing the phagolysosome (PL) and post synaptic density of a synapse onto an encapsulated axon (Ax) (arrows). Scale bar, 300 nm. **h,** 3D rendering of the same branch showing both the phagolysosome (PL) and the axon encapsulated collateral within the OPC cytoplasm. Scale bar, 500 nm. **i-n,** Examples of PLs from different OPCs showing the presence of 40 nm vesicles within the chambers. Scale bar, 300 nm. **o,** 3D reconstruction of an OPC from the P36 dataset. Yellow spheres represent manual annotations of the 189 phagolysosomes found in this OPC. Scale bar, 20 μm (original data in http://www.microns-explorer.org/phagolysosomes/opc). **p,** 3D reconstruction of a microglial cell from the P36 dataset. Red spheres represent manual annotations of the 13 phagolysosomes present in this cell. Scale bar, 20 μm (original data in http://www.microns-explorer.org/phagolysosomes/microglia). **q,** Plot comparing PL density in 9 OPCs and 9 microglia in the P36 dataset.

PLs in OPCs were large enough to visualize their internal contents. Although much of this material was unidentifiable cellular debris, they frequently contained clusters of small (~ 40 nm), clear vesicles (Fig. 2 f,i-n, Extended Data Video 2), suggesting that OPCs had engulfed portions of presynaptic terminals. Occasionally, there was evidence of an entire synaptic terminal surrounded by OPC cytoplasm, with both presynaptic vesicles and a post-synaptic density visible (Fig. 2 f,g). Notably, fragments of myelin, distinct organelles (e.g. mitochondria), and cytoskeletal elements, were not observed in these PLs, suggesting that OPCs primarily engulf axonal processes. In contrast, although microglial PLs also contained unidentifiable electron dense material, they did not contain 40 nm vesicles (Extended Data Fig. 2b-e, h, Video 2), suggesting that, at this age, these two glial cell types recognize and remove distinct cellular elements. The presence of abundant PLs in OPCs containing synaptic material was unexpected, as most structural pruning of neurons is thought to be mediated by microglia^2,28^. To determine the relative abundance of PLs in these two glial cell types, we quantified PLs in complete 3D reconstructions of microglia in the same volumes. This analysis revealed that PLs were significantly more abundant in OPCs than microglia at this age (Fig. 2o,p;Extended Data Fig. 2) (Total PLs/cell: OPCs, 108 ± 48; microglia, 9 ± 3; student’s t-test, p < 0.001, n = 9); although OPCs are larger than microglia, the density of PLs was still significantly higher in OPCs (Fig. 2q; Extended data Fig. 2f,g) (PL density: OPCs, 0.143 ± 0.05 PLs/μm^3^; microglia, 0.03 ± 0.01 PLs/μm^3^; p<0.001, n = 9), suggesting that OPCs are more actively engaged in neuronal process engulfment at this age.

To further explore the prevalence of these organelles in OPCs, OPCs isolated from the visual cortex of one month-old mice were immunolabeled for lysosomal associated membrane protein 2 (LAMP-2), which is associated with both lysosomes and phagolysosomes^29^, and NG2 (chondroitin sulfate proteoglycan 4) that specifically labels OPCs^15^. Consistent with the ultrastructural observations described above, OPCs at this age contained 25 to 54 LAMP-2 immunoreactive circular organelles distributed throughout their somata and processes (Extended Data Fig.1a,b), indicating a high investment in cellular turnover in the developing cortex.

During the critical period of mouse visual cortex from P21 to P35, microglia engulf and eliminate synapses^30,31^ and neuronal corpses through phagocytosis and eventual digestion via acidified phagosomes and PLs s^32^. This structural remodeling of cortical connections declines rapidly after the first postnatal month^2,33^, coinciding with the end of the critical period of ocular dominance plasticity in the visual cortex. To determine if PLs in OPCs follow a similar developmental progression, we quantified PLs in OPCs reconstructed from serial EM volumes obtained in the P49 mouse visual cortex. Significantly fewer PLs were present in OPCs at this age, despite the similar volume occupied by these cells (Extended Data Fig. 3e, f, g) (PL density: P36 OPCs, 0.143 ± 0.05 PLs/μm^3^; P49 OPCs, 0.085 ± 0.03 PLs/μm^3^; p = 0.01, student’s t-test, n = 9, 11, respectively). Together, these results indicate there is a temporal shift in OPC behavior over time and that OPCs may contribute to an early phase of structural remodeling of neurons during the final phase of the critical period in the visual cortex.

OPCs undergo dramatic changes in gene expression and morphology as they differentiate into myelin forming oligodendrocytes^8^ (Extended Data Fig. 3a-d). To determine if axonal engulfment is preferentially associated with the OPC progenitor state, we also examined the cytosol of premyelinating and mature oligodendrocytes at P49 (due to the smaller volume, there was only one partially reconstructed premyelinating oligodendrocyte in the P36 dataset). These more mature oligodendroglia were distinguished from OPCs by the lack of primary cilia^34^, as well as the presence of sheets of membrane extending along axons and membrane wraps indicative of nascent myelination (Extended Data Fig. 3a-d). Premyelinating oligodendrocytes had lower densities of PLs than OPCs PL density: P49 OPCs, 0.085 ± 0.03 PLs/μm^3^; P49 Pre-myelinating oligodendrocyte, 0.034 ± 0.018 PLs/μm^3^; p = 0.001, student’s t-test, n = 11, 5, respectively,(Extended Data Fig. 3f). In fully mature oligodendrocytes, PLs were extremely rare (extended figure 3.e), suggesting that engulfment of neuronal processes declines rapidly as OPCs undergo differentiation (Extended Data, Fig. 3e, f, g).

### OPCs express genes that enable phagocytosis

The 3D EM reconstructions indicate that OPC processes are filled with numerous PLs that contain neuronal material, suggesting that they assist in remodeling nascent neuronal circuits through engulfment of axons and synapses. Phagocytosis requires a complex array of proteins to recognize and engulf distinct cargos, the components for which are highly conserved among different species. To determine whether OPCs express genes that would enable this behavior, we analyzed existing single nucleus RNA-seq and DNA-methylation data sets derived from the mouse motor cortex (P56)^35^ for both lysosomal and phagocytic genes (Fig. 3a,b). OPC nuclei were identified by co-expression of *pdgfra* and *olig2* and premyelinating/myelinating oligodendrocytes were identified by expression of *opalin*(Fig.3 a-c). Expression of genes that encode components of phagocytosis were then compared between OPCs, microglia and mature oligodendrocytes (Fig.3a,b), identified by previous characterization^35^ and expression of distinct marker genes (Extended Data Figure 4).

**Figure 3:**
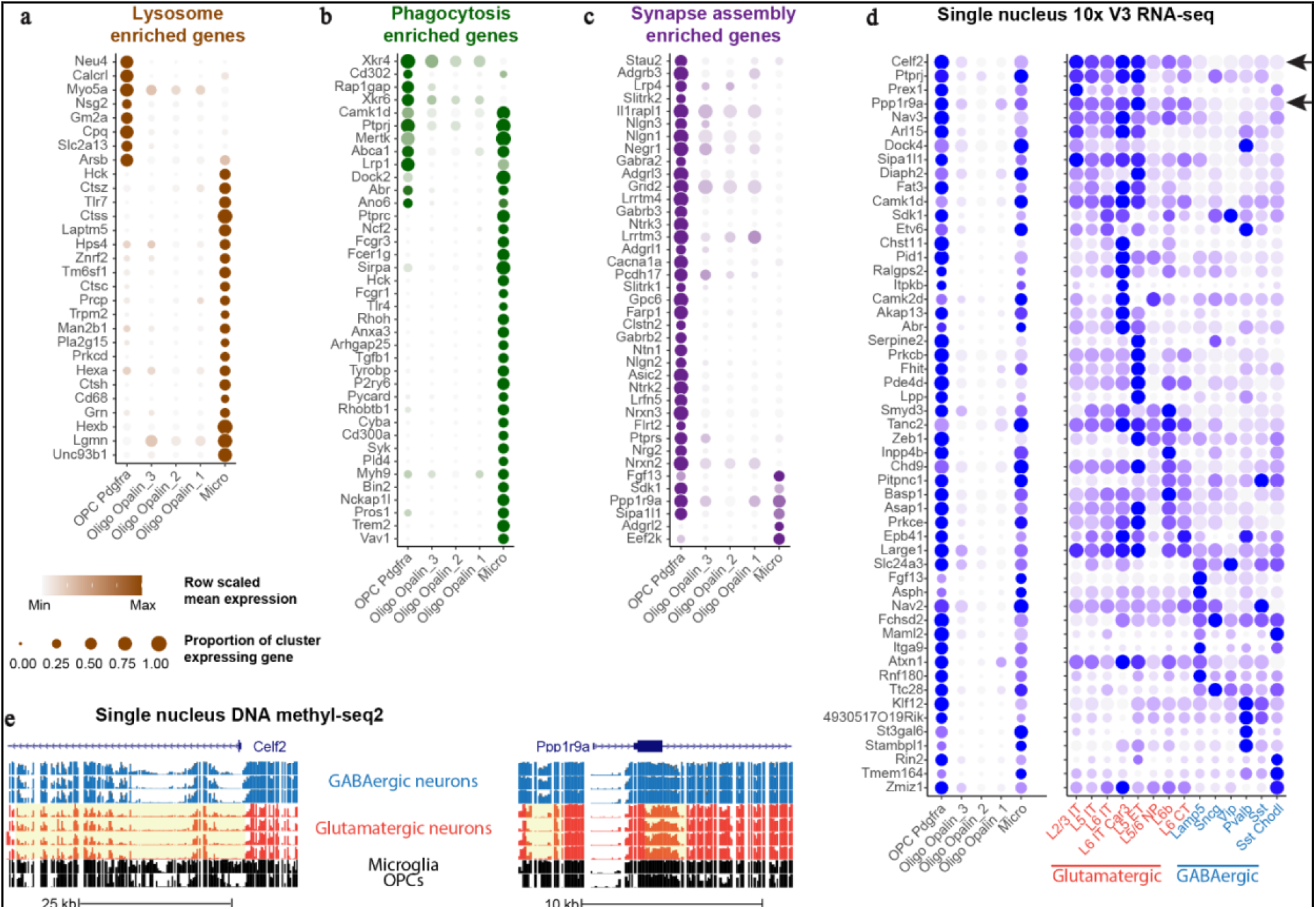
Detection of phagolysosome genes and neuronal transcripts in OPCs, oligodendrocytes and microglia. **a-c**, Dot plots of enriched lysosome (**a**), phagocytosis **(b)**, and synapse assembly **(c)** GO term genes enriched in OPCs and/or microglia, relative to mature oligodendrocytes. **d**, Dot plot of neuronal subclass marker genes expressed in OPCs, mature oligodendrocytes – Oligo, microglia – Micro, and glutamatergic (red) and GABAergic (blue) neuronal subclasses. **e**, Genome tracks showing glutamatergic marker genes, *Celf2* (top panel)_and *Ppp1r9a* (bottom panel) with hypomethylated chromatin highlighted in yellow. Blue tracks show GABAergic and red tracks show glutamatergic neuronal subclasses.

Both OPCs and microglia expressed high levels of mRNAs encoding phagocytic and lysosomal genes. Of the 38 phagocytosis-related genes examined, 32% were expressed in OPCs and 89% were expressed by microglia (Fig.3b). Eight phagocytic genes were expressed in both glial types, including *Mertk, Ptprj*, and *Lrp1. Mertk* (Mer tyrosine kinase) is a member of the TAM (Tyro3-Axl-Mer) family of receptors that signal engulfment, which astrocytes use to refine connectivity in the adult mouse hippocampus by phagocytosing excitatory synapses^36^. *Ptprj* is a tyrosine phosphatase receptor protein that is known to regulate phagocytosis and migration in microglia^37^, and *Lrp1* is a low density lipoprotein receptor essential for myelin phagocytosis and regulation of inflammation in OPCs^38^. Several genes that encode phagocytosis-related proteins expressed in OPCs were absent from microglia: *Rap1gap*, a GTPase activating protein that mediates FcyR dependent phagocytosis^39^; and *Xrkr4* and *Xrkr6*, proteins that promote the exposure of phosphatidylserine to produce “eat me” signaling during phagocytosis, which are highly expressed in the developing brain^40^. The distinct complement of phagocytic genes expressed by OPCs and microglia may enable these cells to identify and engulf discrete parts of neurons. Notably, expression of all the phagocytosis-related genes were negligeable in oligodendrocytes, consistent with the much higher incidence of PLs in OPCs.

We also searched for the presence of neuronal transcripts within microglia and OPCs that may have been engulfed along with neuronal processes. Neuronal transcripts were found in both OPCs and microglia, which suggests both types of glia are phagocytosing neurons at this age Fig.3c,d). However, transcripts encoding these components were higher in OPCs, which correlates with the increased number of PLs in these cells. Because OPCs express many proteins characteristic of neurons, such as AMPA and NMDA glutamate receptors and voltage-dependent Na^+^ channels, we assessed the likelihood that transcription of traditionally “neuronal” genes is occurring directly in OPCs by examining the methylation state of the neuronal genes; genes that are actively transcribed should be hypomethylated. This analysis revealed that microglia and OPCs lack hypomethylation at most neuronal genes (Fig. 3e), a sign that they do not express these genes but incorporated them from elsewhere, likely by phagocytosis of neuronal material. Analysis of cell specific gene expression data suggest that OPCs ingest material from diverse neurons, rather than targeting a specific subtype. These findings, together with the ultrastructural data, provide further evidence that OPCs phagocytose axons of many types of neurons in the mammalian cortex.

### OPCs prune axons in the mouse cortex

The appearance of clusters of 40 nm clear vesicles inside OPC PLs suggests that presynaptic terminals or portions of axons containing terminals are engulfed by these cells. If this process is frequent, OPCs engaged in various stages of this process should be visible in the EM datasets. Indeed, 3D reconstructions revealed that terminal branches of axon collaterals were often surrounded by OPC processes (Fig. 4a-f). The complex anatomy of the branches and the need to accurately manually annotate and quantify the ingestions, made this analysis particularly challenging. Therefore, two individual main branches from two distinct OPCs in the P36 dataset were examined in detail (OPCs #1, #7) (Extended Data Fig. 1c,k; Extended data Fig. 6a-h). We categorized the ingestions as phagosomes (PS), PLs or axon engulfment (Fig. 4g). For the latter, there was no evidence of injury or destruction to the axon, only local contact with OPC cytoplasmic processes that encased the collateral or terminal region of the axon branch. Two types of axon engulfment were observed: either small axonal pieces less than 1 μm in length were surrounded by an OPC process, in which the collateral or terminal axon branch remained connected to the parent axon (24 of 37 in OPC #1, 23 of 32 in OPC #7), or larger axonal collateral or terminal branch segments (up to 3μm) were ensheathed (9 of 37, OPC #1 and 3 of 32 OPC #7), but remained attached to the parent axon (Fig. 4b,e; Extended Data Fig. 4a-h; Video 3).

**Figure 4:**
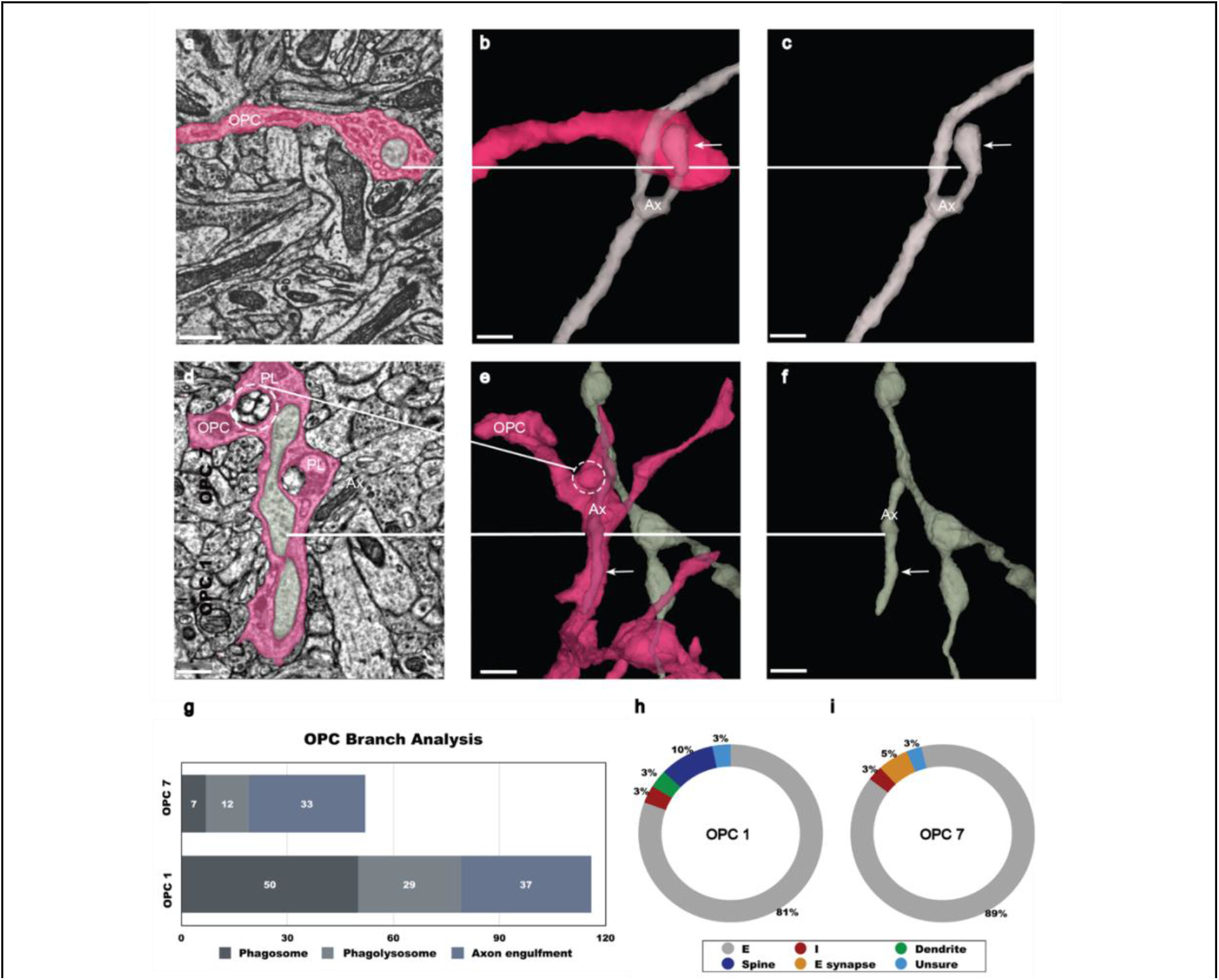
Axon Engulfment by OPCs. **a,** OPC process in pink ingests an excitatory axon bouton (gray) at its tip. Scale bar, 500 nm. **b,** 3D reconstruction reveals a small excitatory axon fragment (Ax) encapsulated within the OPC (pink) at the tip (white line, arrow). Scale bar, 500 nm. **c**, 3D reconstruction shows the same axon (Ax)(arrow) without OPC surrounding it. The bouton tip is visible. Scale bar, 500 nm. **d,** Ultrathin section slice of a large section of an inhibitory axon (Ax)_collateral branch (white line) in gray ingested within the cytoplasm of the OPC (pink). Two phagolysosomes (PL) (dashed circle) are adjacent to the ingested axon. Scale bar, 500 nm. **e,** 3D rendered axon collateral branch encapsulated within the cytoplasm of the OPC (arrow, white line) and phagolysosome within the cytoplasm (dashed circle). Scale bar, 1 μm. **f,** 3D reconstruction of the axon (Ax) in gray shows encapsulated branch (white line, arrow) without the surrounding OPC. Scale bar, 1 μm. **g,** Two individual isolated main branches (one each) of OPC cells #1and #7 were analyzed for ingestion events. Ingestions were categorized as phagosome (PS) phagolysosome (PL) or axon collaterals engulfed in OPC cytoplasm. **h,i,** Donut charts of OPCs 1and 7 quantifies axons partially ingested shows the types-excitatory (E) or inhibitory (I) by examination of boutons and synaptic connections and contact with other neural elements dendrite(D), spine(S), excitatory synapse(Esyn) or unsure (U). Excitatory axons ingestion was most prevalent 89% in OPC#1 and 81% in OPC#7. (See also Extended Data Video 3).

OPCs form synapses with both excitatory and inhibitory neurons in the developing cortex^6,7^ and axons from both neuron classes are targeted for myelination in the cortex^41,42^, indicating that OPCs and their subsequent later developmental stages interact with distinct cell types. To determine if axon pruning is specific to one class of neurons, we traced the branches of 70 axons ensheathed by OPCs back to the main axon and examined the boutons, classifying them as excitatory or inhibitory based on synaptic morphology. This analysis showed the predominant type of axons within the two isolated branches of OPCs were excitatory (89 % in OPC #1 and 81% in OPC #7 of ensheathed structures) (Fig. 4h,i) with 8% undetermined. Approximately 3 % of the engulfed axons were identified as inhibitory, suggesting that OPCs were mainly removing portions of excitatory axons, which comprise the majority of axons in cortical gray matter. This number may be lower than the 10-20% inhibitory neuron population in the neocortex^43^, because the sample size was limited to layer 2/3 and only two OPC branches. Furthermore, the transcriptomics data suggests that OPCs are non-discriminatory phagocytes and engulf all neuron types, based on all layers of the cortex. The association of OPCs with terminal and collateral axon branches was observed in all OPCs from both P36 and P49 datasets, indicating that axon pruning is a conserved function of OPCs during this period. Such associations may represent the predecessor to axonal engulfment and suggest that the terminal portion of axon collaterals are targeted for removal by OPCs.

## Discussion

Consolidation of nascent networks into stable, but adaptable, circuits requires extensive remodeling during development, when excess neurons, axon collaterals and even individual synapses are removed^1,44,45^. This structural refinement is aided by glia, which recognize and engulf both cellular debris and portions of intact neurons^46,47^. Microglia are key participants in this process, as they express phagocytic genes (e.g. *Merktk, Dock2, Sirpa*)^48^, engage in constant surveillance through process motility, can transform into amoeboid, highly phagocytic cells, that accumulate presynaptic material in their cytoplasm^31^. Astrocytes also express phagocytic components (e.g. *Mertk, Megf10*) ^3^ and participate in excitatory synapse removal in adult mice^49^, indicating that structural refinement of neurons is the shared responsibility of multiple glial cell types.

This new analysis of the 3D ultrastructure of individual glial cells in mouse visual cortex using volumetric EM, provides strong evidence that OPCs also participate in neuronal circuit refinement. OPCs, a widely distributed, and abundant population of lineage restricted progenitor cells, extended fine processes that made numerous contacts with terminal axon branches and synapses of excitatory neurons, surrounding and enveloping neuronal components within cup shaped and sheetlike cytoplasmic formations. OPC processes contained a high density of PLs filled with material exhibiting characteristic features of synapses, suggesting that OPCs target specific portions of axons for removal. These ingestions could represent trogocytosis, in which small portions of cells are pinched off^50^. Although the molecular mechanisms that enable trogocytosis have not been fully defined, OPCs expressed numerous genes involved in phagocytosis. In addition, they contained DNA encoding key neuronal constituents, which due to a higher methylation state are likely to have been acquired from neurons. Together, these data provide evidence that OPCs not only serve as progenitors for oligodendrocytes, but also engage in the pruning of neuronal processes during a robust stage of cortical alteration and maturation. By leveraging multiple glial cell types, the nervous system may accelerate circuit refinement by enhancing overall degradative capacity and enable greater spatial and temporal control of this process by recruiting cells that recognize distinct cellular components.

Emerging evidence suggests that OPCs do more than serve as progenitors for oligodendrocytes, as they are found in regions where there is no myelin, and like microglia, OPCs migrate to sites of injury and contribute to scar formation^51–53^. Moreover, recent evidence indicates that OPCs can transform into inflammatory OPCs (*iOPCs*) that engulf and present exogenous antigens through MHC class I and II when exposed to inflammatory cytokines, suggesting that they may modulate tissue inflammation^54,55^. OPCs share many other features with microglia – they are present at a similar density, maintain a grid-like distribution with non-overlapping domains, possess ramified, radially-oriented processes and are highly motile, continuously exploring their surrounding environment with dynamic filopodia^25^.

The robust dynamics and broad distribution of OPCs place them in an ideal position to engage in the modification of neural circuits that we describe in this study. Remarkably, OPC processes contained a higher density of PLs than surrounding microglia, raising the possibility that they are responsible for a greater amount of pruning at this stage of maturation of the cortex. However, it is also possible the engulfment and digestion of axonal elements proceeds more slowly in OPCs than microglia, leading to more PL accumulation. Although the presence of PLs within OPC processes declined with age, the persistence of OPCs throughout the CNS, which retain a similar morphology and dynamics, raises the possibility that they may retain the capacity to modify circuits and clear cellular debris induced by injury or normal aging.

Many OPCs in this region of the developing cortex will differentiate to form oligodendrocytes that myelinate both excitatory and inhibitory axons^43,56–58^. However, the interactions between neurons and OPCs described here are distinct from the early stages of axon wrapping, as these cells had not yet begun transforming into oligodendrocytes, and PL abundance rapidly declined with differentiation. Whether the interactions described here are an early harbinger of myelination^56,59^, to possibly trim superfluous axon branches prior to axon ensheathment or signal which regions of axons are to be myelinated, are not yet known.

These findings raise new questions about how distinct glial cell types coordinate their phagocytic behavior to shape brain connectivity and contribute to plasticity, homeostasis and disease processes. Future studies selectively targeting engulfment by OPCs at different time points may reveal the complex role of these progenitors in early refinement of neural circuits and explore their interactions with astrocytes and microglia.

## Methods

All animal procedures were approved by the Institutional Animal Care and Use Committee at the Allen Institute for Brain Science or Baylor College of Medicine.

### EM Datasets

We used two densely segmented and reconstructed volumes, obtained from serial section electron microscopy (ssEM) in mouse primary visual cortex. These data sets were collected as part of the IARPA Machine Intelligence from Cortical Network (MICrONS) consortium project (https://www.iarpa.gov/index.php/research-programs/microns) to study the functional connectivity of neurons, however they also provide in rich and unprecedented detail the fine morphology of glial cell types. Here we use these datasets to evaluate the population of OPCs and microglia in the 3D environment of the mouse brain visual cortex (preparation methods described below). Dataset 1 was taken from a 200 μm thick sample containing layer 2/3 of mouse visual cortex (P36) and measured approximately 250 μm × 140 μm × 90 μm (Fig.1A). Dataset 2 was taken from a 200 μm thick sample taken from a P49 mouse visual cortex and included all 6 layers, measuring approximately 56 μm × 1 mm × 30 μm. The P36 data set is publicly available at https://microns-explorer.org/.

### Mouse line

Mouse P 36 was a triple-heterozygote for the following three genes: (1) Cre driver: CamKIIa_Cre(Jax:05359<https://www.jax.org/strain/005359>), (2) tTA driver: B6;CBA-Tg(Camk2a-tTA)1Mmay/J (Jax: 003010<https://www.jax.org/strain/003010>), (3) GCaMP6f Reporter: Ai93(JAX 024103) (Allen Institute)^60^. Mouse P49 was a cross of: B6;CBA-Tg(Camk2a-tTA)1Mmay/J (Jax: 003010) and B6;DBA-Tg(tetO-GCaMP6s)2Niell/J (Jax: 024742).

### Two-Photon Imaging

Before preparation for electron microscopy, mice underwent neurophysiology data acquisition conducted at Baylor College of Medicine (details in ^9^). Note that this 2-photon data was not used in this study. Briefly, a 3 mm craniotomy was made centered on the primary visual cortex (V1; 2.7mm lateral of the midline, contacting the lambda suture), and the cortical window was then sealed with a 3 mm coverslip (Warner Instruments), using cyanoacrylate glue (VetBond). The mouse was allowed to recover for 1-2 hours prior to the imaging session. Imaging was performed in V1, in a 400 × 400 × 200 μm3 volume with the superficial surface of the volume at the border of L1 and L2/3, approximately 100um below the pia. Laser excitation was at 920nm at 25-45mW depending on depth. The objective used was a 25x Nikon objective with a numerical aperture of 1.1, and the imaging point-spread function was measured with 500 nm beads and was approximately 0.5 × 0.5 × 3 μm3 in x, y, and z. To aid in registration of optical physiology data to EM data, a wide field image of the cranial window visualizing the surface vasculature was provided in addition to a volumetric image stack of the vasculature, encompassing the region of tissue where the neurophysiology dataset was acquired. The vasculature was imaged by subcutaneous injection of 60 μL 2.5% Texas Red 3000MW Lysine fixable (Life Technologies D3328, allowing blood vessels and GCaMP6-expressing cell bodies to be imaged simultaneously by 2-photon microscopy. Mice were then transferred to the Allen Institute in Seattle and kept in a quarantine facility for 1 to 3 days, prior to perfusion.

### Perfusion

After induction of anesthesia with isoflurane, the appropriate plane of anesthesia was checked by a lack of toe pinch reflex and the animals were transcardially perfused with 15 ml 0.15 M cacodylate buffer (EMS, Hatfield, PA, pH 7.4) followed by 30 ml fixative mixture containing 0.08 M cacodylate (pH 7.4), 2.5% paraformaldehyde (EMS), 1.25% glutaraldehyde (EMS) and 2 mM calcium chloride (Sigma). The perfusion solution was based on the work of (Hua et al., 2015). Once the brain was removed it was placed into the same fixative solution to post-fix for 16 to 72 hours at 4 °C.

After perfusion of the animals and excision of the brain, the surface of the cortex was imaged using differential contrast lighting to visualize the surface vasculature of visual cortex and identify the region where the cranial window had previously been. This region was then prepared for Electron Microscopy. Details of the procedures to carefully map the neurophysiology site in the histological sections is described in (details in ^9^).We omitted these details here as no neurophysiology data was used in this manuscript, even though the anatomical data originates from the same block of tissue that was recorded with two-photon imaging. The brain was washed in CB (0.1 M cacodylate buffer pH 7.4) and embedded in 2% agarose. The agarose was trimmed and mounted for coronal sectioning in a Leica VT1000S vibratome; successive 200 μm thick slices were taken until the entire region of cortical tissue previously demarcated by manual markings was sectioned. During this procedure, we also acquired blockface images of each brain slice. The coronal sections containing the imaged site were then selected for histological processing (see below).

### EM Histology

The histology protocol used here is based on the work of ^61^ and ^62^, with modifications to accommodate different tissue block sizes and to improve tissue contrast for transmission electron microscopy (TEM). Following several washes in CB (0.1 M cacodylate buffer pH 7.4), the vibratome slices were treated with a heavy metal staining protocol. Initial osmium fixation with 2% osmium tetroxide in CB for 90 minutes at room temperature was followed by immersion in 2.5% potassium ferricyanide in CB for 90 minutes at room temperature. After 2 × 30 minute washes with deionized (DI) water, the tissue was treated with freshly made and filtered 1% aqueous thiocarbohydrazide at 40 °C for 10 minutes. The samples were washed 2 × 30 minutes with DI water and treated again with 2% osmium tetroxide in water for 30 minutes at room temperature. Double washes in DI water for 30 min each were followed by immersion in 1% aqueous uranyl acetate overnight at 4°. The next morning, the samples in the same solution were placed in a heat block to raise the temperature to 50° for 2 hours. The samples were washed twice in DI water for 30 minutes each, then incubated in Walton’s lead aspartate pH 5.0 for 2 hours at 50 °C in the heat block. After double washes in DI water for 30 minutes each, the slices were dehydrated in an ascending ethanol series (50%, 70%, 90%, 3 × 100%) 10 minutes each and two transition fluid steps of 100 % acetonitrile for 20 minutes each. Infiltration with acetonitrile:resin dilutions at 2p:1p (24 h), 1p:1p (48 h) and 1p:2p (24 h) were performed on a gyratory shaker. Samples were placed in 100% resin for 24 hours, followed by embedment in Hard Plus resin (EMS, Hatfield, PA). The samples were cured in a 60 °C oven for 96 hours.

In order to evaluate the quality of samples during protocol development and before preparation for large scale sectioning, the following procedure was used for tissue mounting, sectioning and imaging. We evaluated each sample for membrane integrity, overall contrast and quality of ultrastructure. For general tissue evaluation, adjacent slices and tissue sections from the opposite hemisphere, processed in the same manner as the ROI slice, were cross-sectioned and thin sections were taken for evaluation of staining throughout the block neighboring the region of interest.

### Ultrathin Sectioning

The tissue block was trimmed to contain the neurophysiology recording site which is the region of interest (ROI) then sectioned to 40 nm ultrathin sections. For both trimming and sectioning a Leica EM UC7 ultramicrotome was equipped with a diamond trimming tool and an Ultra 35 diamond knife (Diatome USA) respectively. Sectioning speed was set to 0.3 mm/sec. Eight to ten serial thin sections were cut to form a ribbon, after which the microtome thickness setting was changed to 0 nm in order to release the ribbon from the knife edge. Then, using an eyelash superglued to a handle, ribbons were organized to pairs and picked up as pairs to copper grids (Pelco, SynapTek, 1.5 mm slot hole) covered by 50nm thick LUXFilm support (Luxel Corp., Friday Harbor, WA).

### Electron microscopy imaging

The imaging platform used for high throughput serial section imaging is a JEOL-1200EXII 120kV transmission electron microscope that has been modified with an extended column, a custom scintillator, and a large format sCMOS camera outfitted with a low distortion lens. The column extension and scintillator facilitate an estimated 10-fold magnification of the nominal field of view with negligible impact on resolution. Subsequent imaging of the scintillator with a high-resolution, large-format camera allows the capture of fields-of-view as large as 13×13 μm at 4 nm resolution. As with any magnification process, the electron density at the phosphor drops off as the column is extended. To mitigate the impact of reduced electron density on image quality (shot noise), a high-sensitivity sCMOS camera was selected and the scintillator composition tuned in order to generate high quality EM images within exposure times of 90 - 200 ms^63^.

### Image volume assembly and morphological segmentation

Aligning the individual image tiles and sections into a coherent three-dimensional volume and segmenting the cellular morphology for the P36 dataset was performed as previously described within^9–11^.

### Proofreading and Annotation of Volumetric Imagery Data

We used a combination of Neuroglancer (Maitin-Shepard, https://github.com/google/neuroglancer) and custom tools to annotate and store labeled spatial points^64^. In brief, we used Neuroglancer to simultaneously visualize the imagery and segmentation of the 3d EM data. A custom branch of Neuroglancer was developed that could interface with a “dynamic” segmentation database, allowing users to correct errors (i.e., either merging or splitting neurons) in a centralized database from a web browser. Neuroglancer has some annotation functionality, allowing users to place simple annotations during a session, but does not offer a way to store them in a central location for analysis. We thus built a custom cloud-based database system to store arbitrary annotation data centered associated with spatial points that could be propagated dynamically across proofreading events. Annotations were programmatically added to the database using a custom python client and, in relevant cases, after parsing temporary Neuroglancer session states using custom python scripts. These spatial points and their associated data (e.g., synapse type, cell body ID number, or cell types) were linked to stored snapshots of the proofreading for querying and reproducible data analysis. All data analyzed here came from the “v183” snapshot.

### Visualization and Analysis of Mesh Data

Neuronal meshes were computed by Igneous (https://github.com/seung-lab/igneous) and kept up to date across proofreading. Meshes were analyzed in a custom python library, MeshParty (https://github.com/sdorkenw/MeshParty), that extends Trimesh (https://trimsh.org) with domain-specific features and VTK (https://www.vtk.org) integration for visualization. In cases where skeletons were used, we computed them with a custom modification of the TEASAR algorithm^65^ on the vertex adjacency graph of the mesh object implemented as part of MeshParty. In order to associate annotations such as synapses or AIS boundary points with a mesh, we mapped point annotations to the closest mesh vertex after removing artifacts from the meshing process.

### Quantification and Statistical Analysis

A t-test (two sample assuming unequal variance, two tailed) was performed in Microsoft Excel to compare number and densities of phagolysosomes.

### Annotation of cells in the EM volume

We used a combination of Neuroglancer (Maitin-Shepard https://github.com/google/neuroglancer) and custom tools to annotate and store labeled spatial points. In brief, we used Neuroglancer to simultaneously visualize the imagery and segmentation of the 3d EM data. Neuroglancer incorporates the capability to store and remark on xyz points within the data. These spatial points and their associated data (e.g., Cell type, cell body ID number, or organelles) were linked to stored spreadsheets and documents.

### Mouse primary OPC culture

Cerebral cortices from post-natal day 6 or 7 (P6/P7) CD1 mouse pups (Charles River) were collected, and tissue was dissociated using MACS Milltenyi Biotec Neural Tissue Dissociation Kit (P) (130-092-628), according to the manufacturer’s instructions. Briefly, cortices were collected from each pup and incubated at 37°C with gentle rotation using a MACSmix^TM^ Tube Rotator (130-090-753) following digestion with pre-warmed enzyme P mix. Cells were then mechanically dissociated and passed through a 70μm MACS SmartStrainer and a 70μm Pre-Separation Filter to remove cell clumps before incubation with Anti-O4 Microbeads (130-094-543). Cells were then loaded into an MS column (130-042-201) in for magnetic separation, and positively-selected cells were collected following extrusion through the column.

Cells were plated at a density of about 20,000 cells / well in a 24-well plate with 1.5mm coverslips in each well that had previously been coated overnight with poly-d-lysine (50μL PDL in 250mL dH_2_O). Cells were expanded overnight at 37°C in OPC media containing recombinant human PDGF-AA and Neurotrophin-3 until optimal density was reached. The plates were then processed for immunofluorescence staining.

### Staining of primary OPC cultures

Coverslips were washed once with 1x PBS before 4% PFA was applied for 13 minutes at room temperature. Cells were then washed twice with 1x PBS (5 minutes/wash) before blocking in 0.5% Saponin, 5% normal donkey serum, and 1x PBS) for 1 hour at room temperature. Cells were then stained with primary antibodies against NG2 proteoglycan (guinea pig, 1:100, custom Bergles antibody), LAMP2 (rat, Abcam, 1:200) to visualize OPC cell bodies/ processes and lysosomes, respectively. Following overnight incubation at 4°C with rotation, cells were washed three times with 1x PBS (5 minutes/wash) at room temperature. AlexaFluor secondary antibodies (488 donkey anti-rat and 647 donkey anti-guinea pig) were applied for 1 hour at room temperature in the dark. Cells were again washed three times, DAPI stain was applied for 10 minutes, and cells were washed again twice before coverslips were removed and mounted to slides using ProLong^TM^ Gold Antifade Mountant and allowed to dry at room temperature for 24 hours.

### Imaging and analysis of primary OPC cultures

Cells from three separate culture experiments were used for the analysis. All images were taken at 63x magnification using the AiryScan function on a Zeiss 710 confocal microscope. Airyscan-processed z-stacks of NG2^+^ individual OPCs were projected using the maximum intensity projection function, and the number of LAMP2^+^ vesicles within cell processes were quantified.

### Reagents used in primary OPC cultures

*OPC Culture Tools* – **all Miltenyi** (Cat no.)

MACS Milltenyi Biotec Neural Tissue Dissociation Kit (P) (130-092-628)

MACS Milltenyi Biotec Anti-O4 MicroBeads (130-094-543)

MACS SmartStrainer (70μm) (130-090-753)

MACSmix^TM^ Tube Rotator (130-090-753)

Pre-Separation Filters, 70μm (130-095-823)

MACS MS Columns (130-042-201)

MACS Multistand (130-042-303)

MiniMACS^TM^ Separator (130-042-102)

### Antibodies

Anti-NG2 (Guinea pig, source: Bergles Laboratory)

Anti-LAMP2 (Abcam, cat no. ab13524)

### Additional Reagent

0.5% saponin (Millipore Sigma, 47036)

ProLong^TM^ Gold Antifade Mountant (ThermoFisher, cat no. P10144)

### Experimental Models: Organisms/ Strains

CD1 mice, Charles River (Strain number 022)

### Molecular analysis of OPCs and microglia with single nucleus RNA-seq and single nucleus DNA methyl-seq2 dataset

We analyzed recently described Chromium 10x V3 single nucleus RNA-seq and single nucleus DNA methyl-seq2 datasets from mouse primary motor cortex^35^, available from the Neuroscience Multi-omic Data Archive (NeMO, https://assets.nemoarchive.org/dat-ch1nqb7). Only the 159,738 nuclei from the dataset generated by the Broad Institute were used.

For gene expression dot plots, the mean UMIs per cluster and proportion of cluster with UMIs greater than 1 were calculated using the scrattch.hicat v0.0.22 library (https://github.com/AllenInstitute/scrattch.hicat) and visualized using ggplots2 v3.3.3 library^66^ in R v3.4.1. Mean expression was scaled from 0 to 1 for each gene for each major cell class (i.e. non-neuronal and neurons). To find neuronal subclass marker genes, we first created a Seurat ^67, 68^ object of only the neuronal cell types, downsampled to 500 nuclei per neuronal subclass. We then used the FindAllMarkers function from Seurat v3.2.0 with the ‘roc’ test to identify differentially expressed genes that were enriched in a particular neuronal subclass compared to other neurons. Neuronal subclass marker genes were then filtered to include genes with greater than 2 log2FC in OPCs and microglia relative to each mature oligodendrocyte cluster (Oligo Opalin_1-3), and with greater than 2 log2 cpm expression in OPCs and microglia.

To visualize single nucleus DNA methyl-seq2 tracks, we loaded gene body methylation (CGN) tracks into the UCSC genome browser. Example tracks of neuronal subclass marker genes that showed expression in OPCs were identified to highlight the lack of DNA hypomethylation in OPCs.

Specific GO term gene lists for lysosome, phagocytosis, and synapse assembly were downloaded from http://www.informatics.jax.org/go/term/ on January 20^th^, 2021^69,70,71^. Each gene list was filtered to include genes with greater than 2 log2FC in OPCs and/or microglia relative to each mature oligodendrocyte cluster, and with greater than 2 log2 cpm expression in OPCs and/or microglia.

## Data availability

The raw images, segmentation, and synaptic connectivity will be made available upon or before publication.

## Code availability

All software is open source and available at http://github.com/seung-lab if not otherwise mentioned.

Alembic: Stitching and alignment.

CloudVolume: Reading and writing volumetric data, meshes, and skeletons to and from the cloud

Chunkflow: Running convolutional nets on large datasets

DeepEM: Training convolutional nets to detect neuronal boundaries.

DynamicAnnotationFramework: Proofreading and connectome updates (visit https://github.com/seung-lab/AnnotationPipelineOverview for repository list)

Igneous: Coordinating downsampling, meshing, and data management.

MeshParty: Interaction with meshes and mesh-based skeletonization (https://github.com/sdorkenw/MeshParty)

MMAAPP: Watershed, size-dependent single linkage clustering, and mean affinity agglomeration.

PyTorchUtils: Training convolutional nets for synapse detection and partner assignment (https://github.com/nicholasturner1/PyTorchUtils).

Synaptor: Processing output of the convolutional net for predicting synaptic clefts (https://github.com/nicholasturner1/Synaptor).

TinyBrain and zmesh: Downsampling and meshing (precursors of the libraries that were used).

## Author Contributions

Conceptualization J.B., D.E.B., N.L. J., T.E. B., N.M.C; Software (Alignment) G.M., F.C., R.T., T.M.,W.M.S and W.W.; Software (Analysis infrastructure) C.S.J., C.S.M., F.C and S.D., S.S..; Software (Data interaction and viewing) M.C., N.K. and W.M.S.; Software (Cloud data storage) W.M.S.; Software Proofreading System) J.Zu., N.K. and S.D.; Formal Analysis A.N., B.H., C.S.M., F.C., J.Zh. and S.D.; Investigation (Histological preparation for EM), J.B., M.T. and N.M.C.; Investigation (Sectioning sample for EM), A.A.B and A.L.B, D. J. B..; Investigation (EM Imaging), W. Y., D.B., A.A.B, A.L.B, D.J.B, N.M.C and R. C. R..; Investigation (Alignment) D.I., G. M., I.T., R.T, T.M., Y. L. and W.W.; Investigation (Neuron Reconstruction) A.Z., I.T., J. Zu., J.W., K.L., M.C., N.K., R.L., S.P., S. M. and T.M.; Investigation (Synapse Detection) J.W and N.L.T.; Investigation (Cell Typing): J.B, L. E., F.C., C.O., A.M.W. and N.M.C.; Data Curation (Proofreading) N.M.C., Investigation (Sub cellular structures): C. O., J.B. and J.L.-S Investigation (Genomics): N.L. J., T.E. B, R. H. and E.S. L. Investigation (OPC cultures): J. G. and D. E. B. Writing-original J.B., D.E.B., N.L. J., T.E. B., N.M.C. Writing – Review & Editing F.C., H.S.S., R. C.R., C. O., M. T., N. T.; Visualization C.S.M. and F.C., Supervision E.S. L., J.L.-S., D.E. B., H. S.S., N.M.C. and R.C.R., Project administration H.S.S., N.M.C. and R.C.R., Funding Acquisition H.S.S., N.M.C. and R.C.R.

## Acknowledgments

We thank John Philips, Sill Coulter and the Program Management team at the AIBS for their guidance for project strategy and operations. We thank Hongkui Zeng, Christof Koch and Allan Jones for their support and leadership. We thank Andreas Tolias, Jacob Reimer and their teams at the Baylor College of Medicine for providing the mice used for electron microscopy. We thank the Manufacturing and Processing Engineering team at the AIBS for their help in implementing the EM imaging and sectioning pipeline. We thank Rob Young and the Technology team at the AIBS for their work in the imaging processing of the EM data. We thank Brian Youngstrom, Stuart Kendrick and the Allen Institute IT team for support with infrastructure, data management and data transfer. We thank the Facilities, Finance, and Legal teams at the AIBS for their support on the MICrONS contract. We thank Stephan Saalfeld for help with the parameters for 2D stitching and rough alignment of the dataset. We would like to thank the “Connectomics at Google” team for developing Neuroglancer and computational resource donations. We also would like to thank Amazon and Intel for their assistance. We thank S. Koolman, M. Moore, S. Morejohn, B. Silverman, K. Willie, and R. Willie for their image analyses, Garrett McGrath for computer system administration, and May Husseini and Larry and Janet Jackel for project administration. We thank Kristina Micheva and Stephen J Smith for advice and feedback. We thank Prof Joseph Ayers, Erin Cram and James Monaghan for guidance, support and feedback. Supported by the Intelligence Advanced Research Projects Activity (IARPA) via Department of Interior/ Interior Business Center (DoI/IBC) contract numbers D16PC00003, D16PC00004, and D16PC0005. The U.S. Government is authorized to reproduce and distribute reprints for Governmental purposes notwithstanding any copyright annotation thereon. Disclaimer: The views and conclusions contained herein are those of the authors and should not be interpreted as necessarily representing the official policies or endorsements, either expressed or implied, of IARPA, DoI/IBC, or the U.S. Government. HSS also acknowledges support from NIH/NINDS U19 NS104648,and research reported in this publication was supported by the National Institute of Mental Health of the National Institutes of Health under Award Number U19MH114824.

ARO W911NF-12-1-0594, NIH/NEI R01 EY027036, NIH/NIMH U01 MH114824, NIH/NINDS R01NS104926, NIH/NIMH RF1MH117815. NIH/NIA R01AG072305 (DB), the Dr. Miriam and Sheldon G Adelson Medical Research Foundation (DB), and the Mathers Foundation. The content is solely the responsibility of the authors and does not necessarily represent the official views of the National Institutes of Health.

We thank the Allen Institute for Brain Science founder, Paul G. Allen, for his vision, encouragement and support.

## Extended data figures

**Extended data Figure 1.**
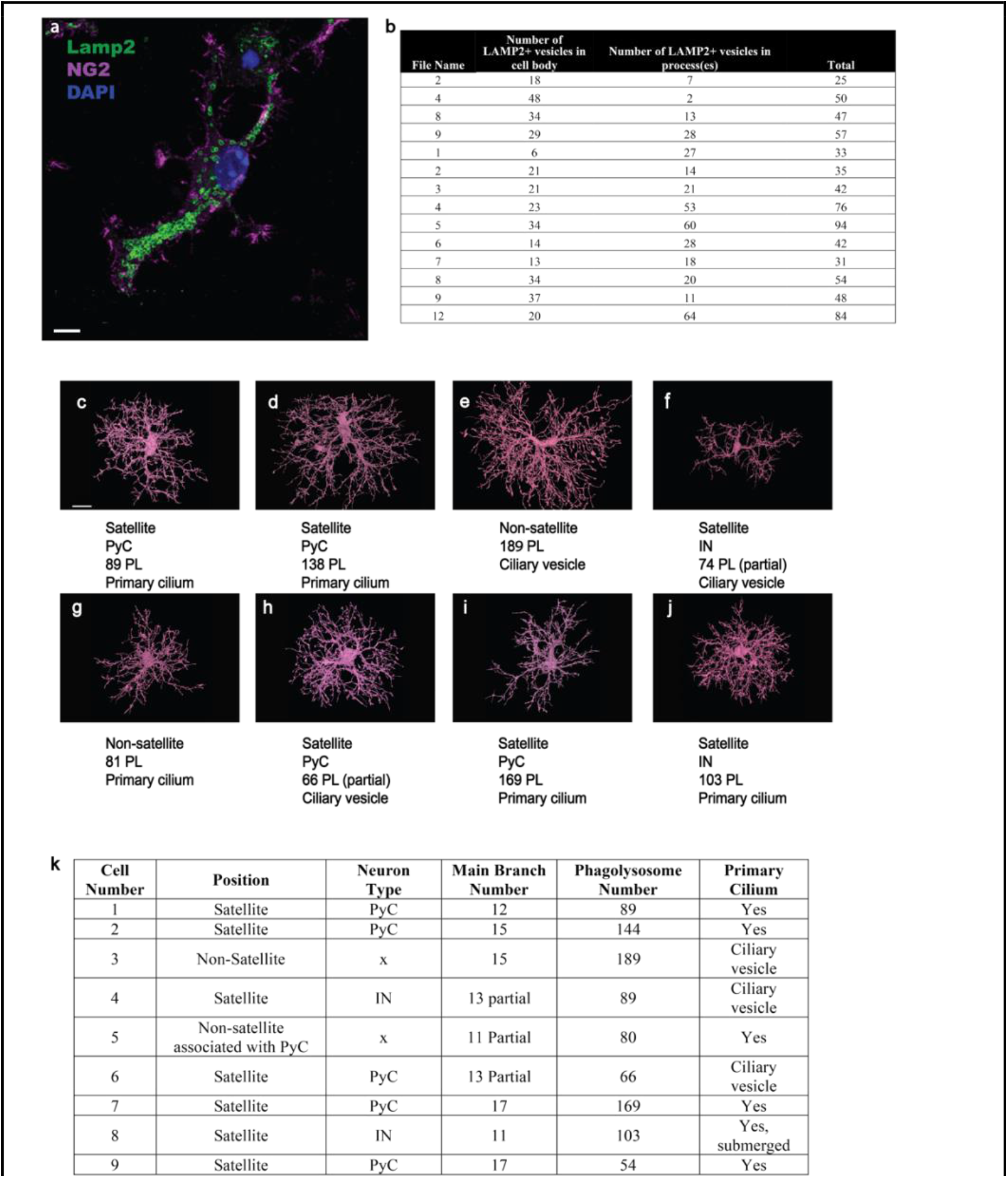
Phagolysosome counts and OPC features. **a,** Primary cultured OPC, immunolabeled with Lamp2 (green)and NG2 chondroitin-sulfate proteoglycan (fuchsia), nucleus stained in blue (DAPI). Scale bar 1μm. **b,** Quantification of Lamp2 positive organelles (lysosomes and phagolysosomes) in OPC soma and branches. **c-j**. Vignettes of 8 of 16 OPCs used for analysis in the P36 dataset. the ninth cell is pictured in Fig. 2a. Scale bar for all cells, 15 μm. **k,** Cells were scored for satellite position, neuron host type, number of main branches, number of phagolysosomes and presence of primary cilia.

**Extended Data Figure 2.**
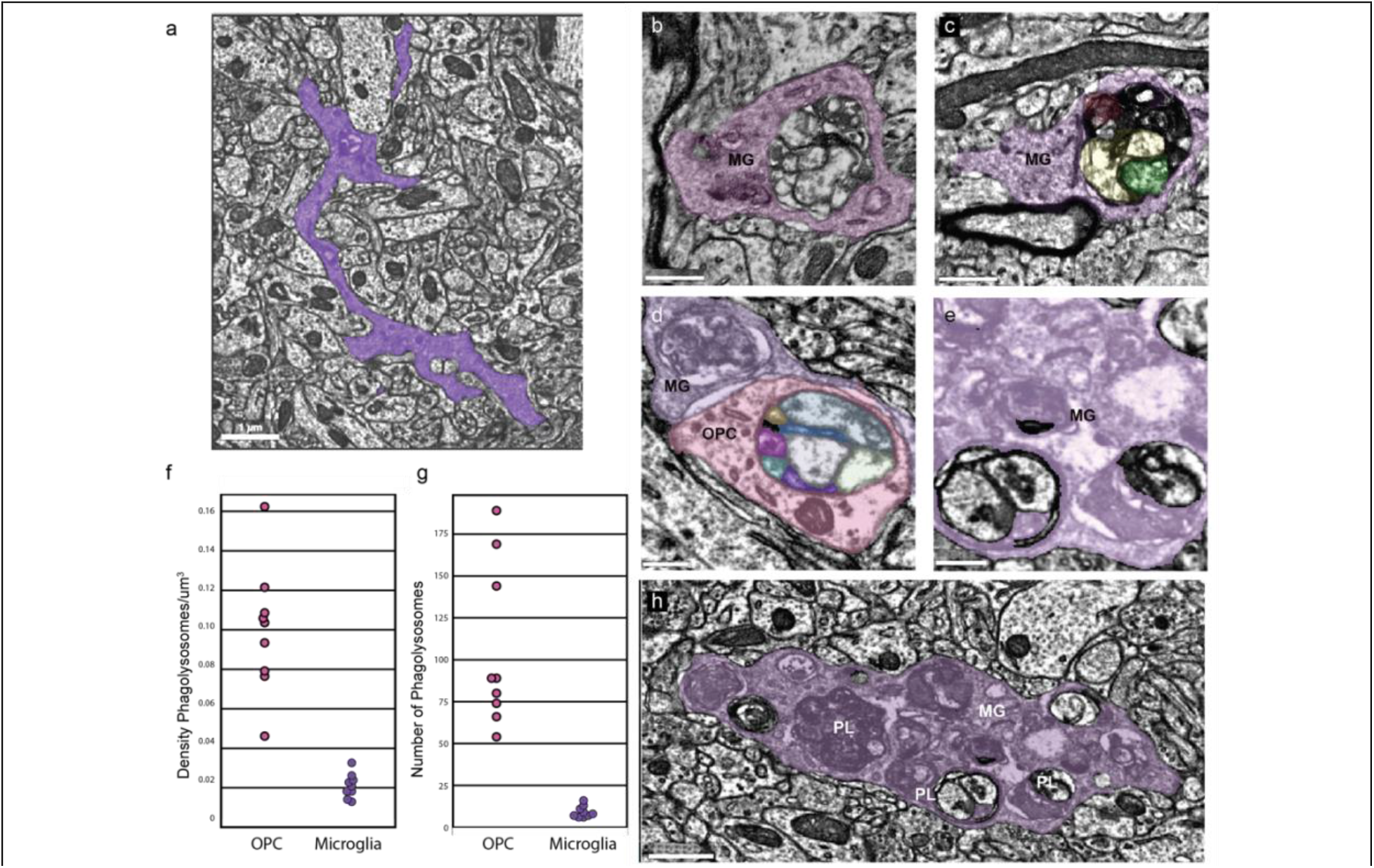
Phagolysosomes in Microglia. **a,** Microglial cell branch in purple shows a phagolysosome (arrow) in its dense cytoplasm. Scale bar, 1 μm. **b,** and **c,** Two large phagolysosomes from P49 dataset microglial cells (MG) in purple reveal electron dense material ingested within are in close proximity to myelinated axons (arrows). Scale bars, 500 nm. **d,** A microglial (MG) branch (purple) from P36 dataset contacts and OPC branch in pink with a phagolysosomes in both cells. Scale bar, 300 nm. **e,** A cluster of phagolysosomes in a microglial cell from P36 dataset shows electron dense material within without any sign of 40 nm vesicles. Scale bar, 300 nm. **f,** Comparison of density of phagolysosomes at P36 between OPCs and microglia (density was calculated including the volume of the soma region) **g,** Comparison of the total number of phagolysosomes at P36 between OPCs and microglia **h,** A lower magnification view of the branch in e shows the cytoplasm congested with phagolysosomes(PL). Scale bar, 750 nm. **f, g,** Plot shows the comparison of phagolysosome numbers in the microglia in P36 and P49 datasets.

**Extended data Figure 3.**
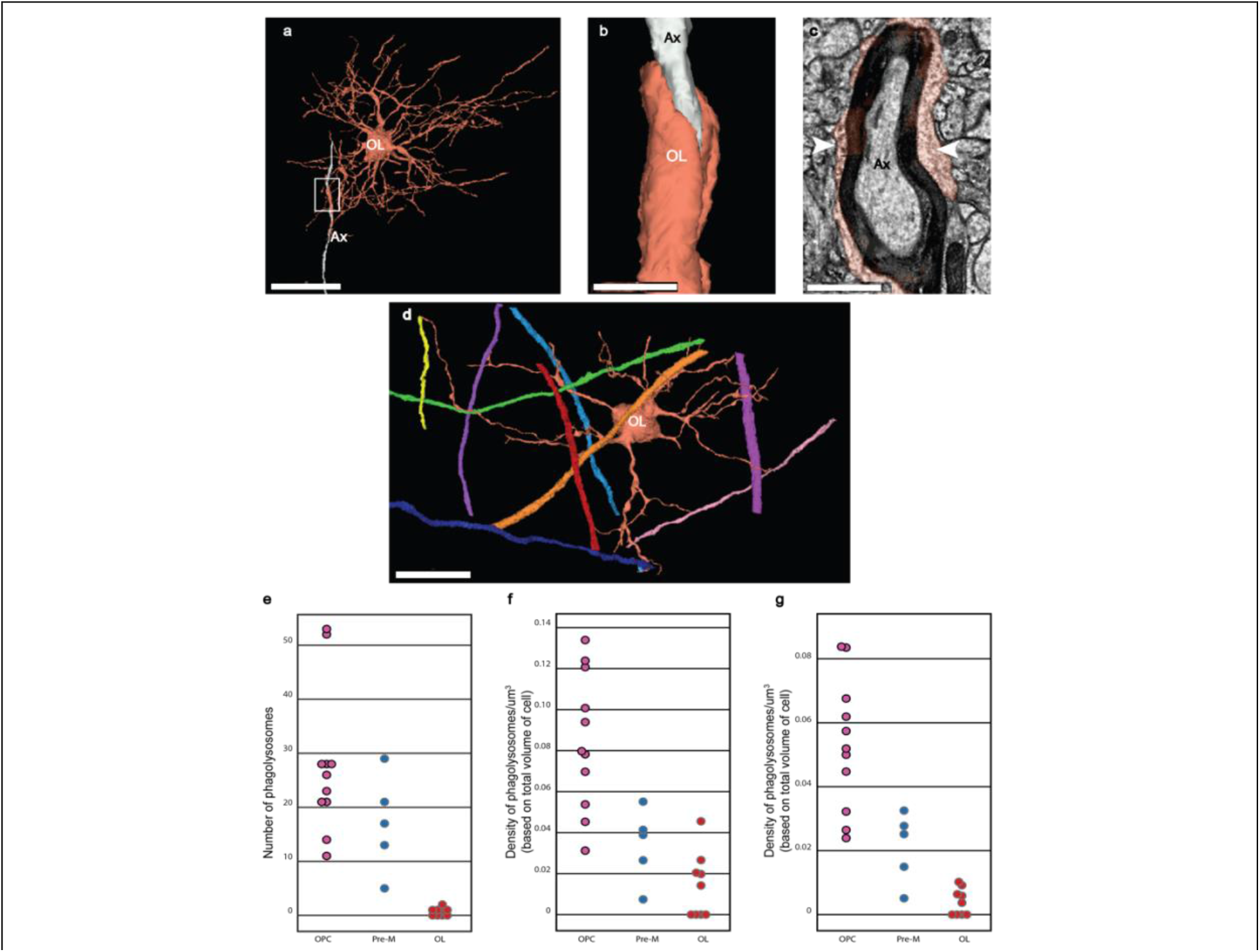
Premyelinating cells, myelinating oligodendrocyte and phagolysosome densities. **a,** 3D rendering of a premyelinating oligodendrocyte (OL) in P49 dataset shows a branch aligned with an axon (white)(boxed area). Scale bar, 20 μm. **b,** Higher magnification view of boxed area shows myelin (orange) and axon (white). Scale bar, 1.5 μm. **c,** Ultrathin section view shows the uncompacted myelin(orange)(arrows) on the outside of the compact myelin (black). Scale bar, 750 nm. **d,** Mature myelinating oligodendrocyte (OL) has thin branches that are aligned with colored myelinated axons. Scale bar, 10μm. **e-g,** Plots comparing densities of phagolysosome distribution in OPCs, Premyelinating cells and mature oligodendrocytes.

**Extended Data Figure 4.**
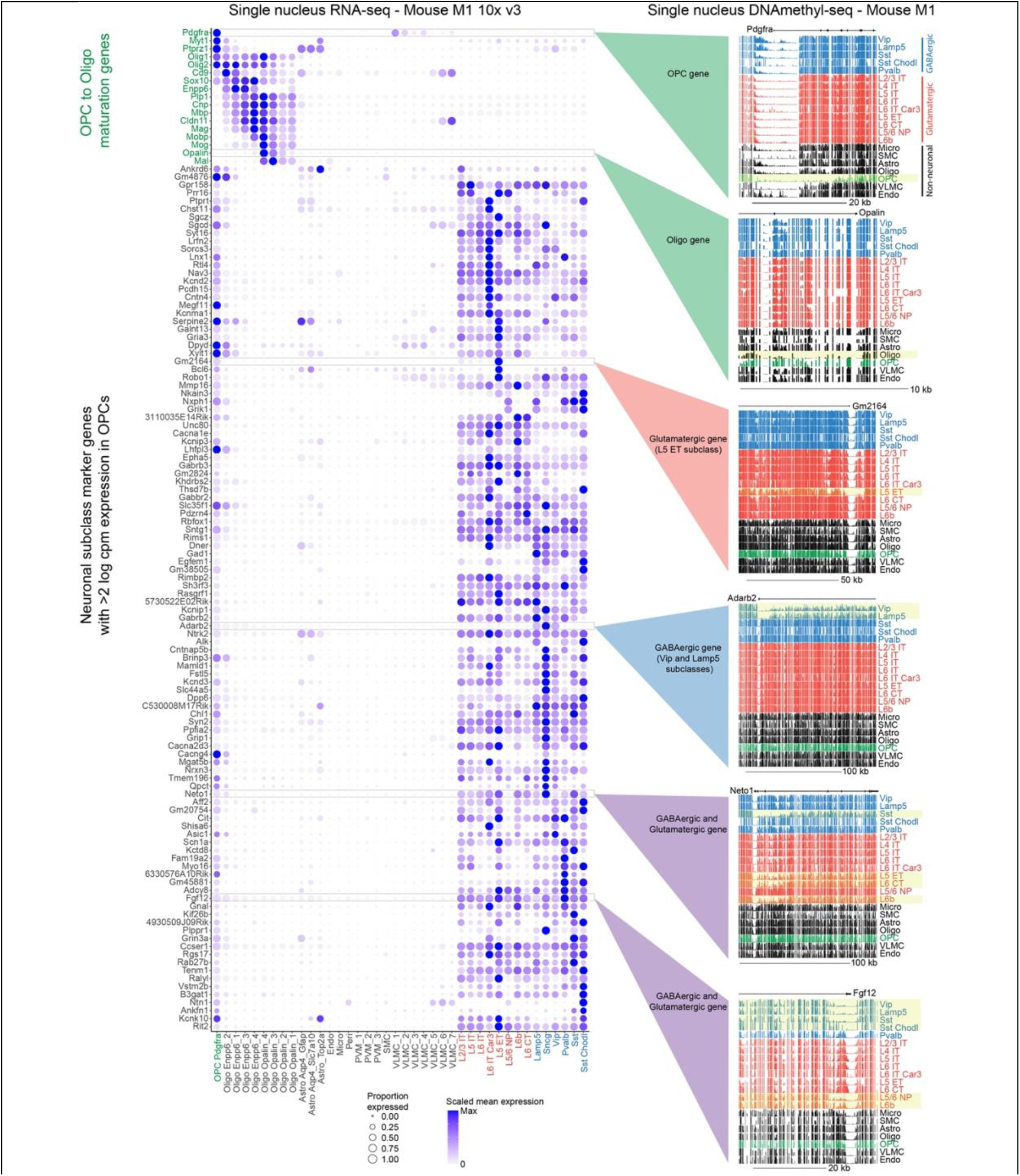
Transcriptomic and epigenomic profiling of neuronal subclass genes in OPCs. Dot plot of OPC to oligodendrocyte maturation genes (green) and neuronal subclass marker genes for all non-neuronal clusters and neuronal subclasses in dataset. Expression is scaled by gene across all clusters and subclasses. Blue indicates GABAergic and red indicates glutamatergic neuronal subclasses. Clusters with of gene body DNA hypomethylation are highlighted in yellow.

**Extended Figure 5.**
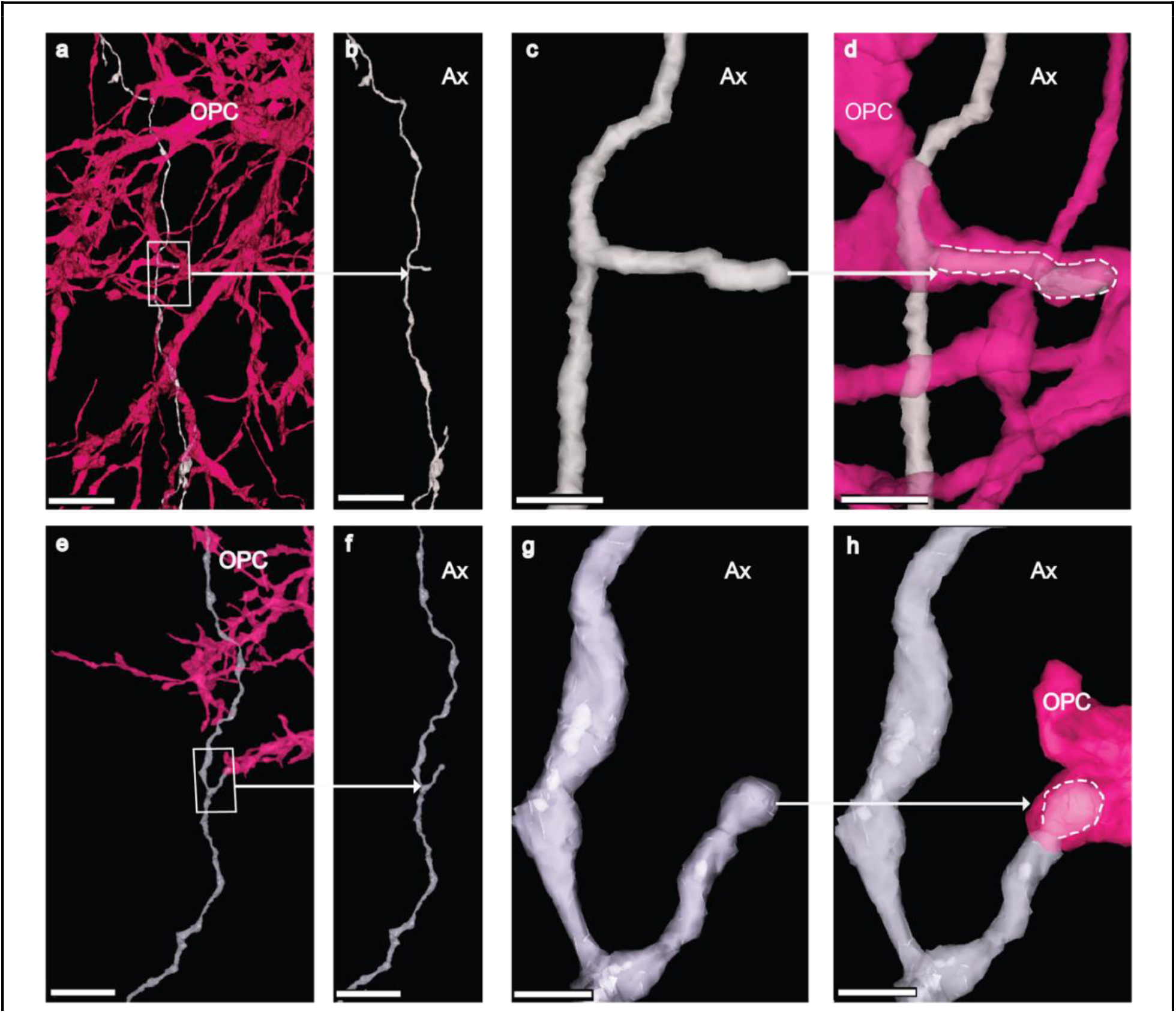
OPCs ingest collateral branches. **a**, OPC (pink) contacts an excitatory axon (gray). Boxed area shows site of ingestion. Scale bar, 5 μm. **b**, 3 D rendering of the excitatory axon (gray). Arrow points to collateral branch in boxed area in (**a)**. Scale bar, 5 μm. **c**, Higher magnification of collateral branch. White arrow points to ingested branch in (**d)**. Scale bar, 750 nm. **d,** 3D rendering of excitatory axon (gray) ingested within the branch of an OPC (pink). Dotted line indicates the outline of ingested collateral branch. Scale bar, 750 nm. **e**, OPC (pink) contacts inhibitory axon (gray). Boxed area shows site of ingestion. Scale bar, 5 μm. **f**, 3D rendering of inhibitory axon (gray). Arrow points to collateral branch in boxed area in (**e**). Scale bar, 5 μm. **g,** Higher magnification of collateral branch. White arrow points to ingested branch in (**e)**. Scale bar, 750 nm. **h,** 3D rendering of inhibitory axon (grey) ingested within the branch of an OPC (pink). Dotted line indicates the outline of ingested collateral branch. Scale bar, 750 nm. Also see Extended Data Video 3.

**Extended Data Figure 6.**
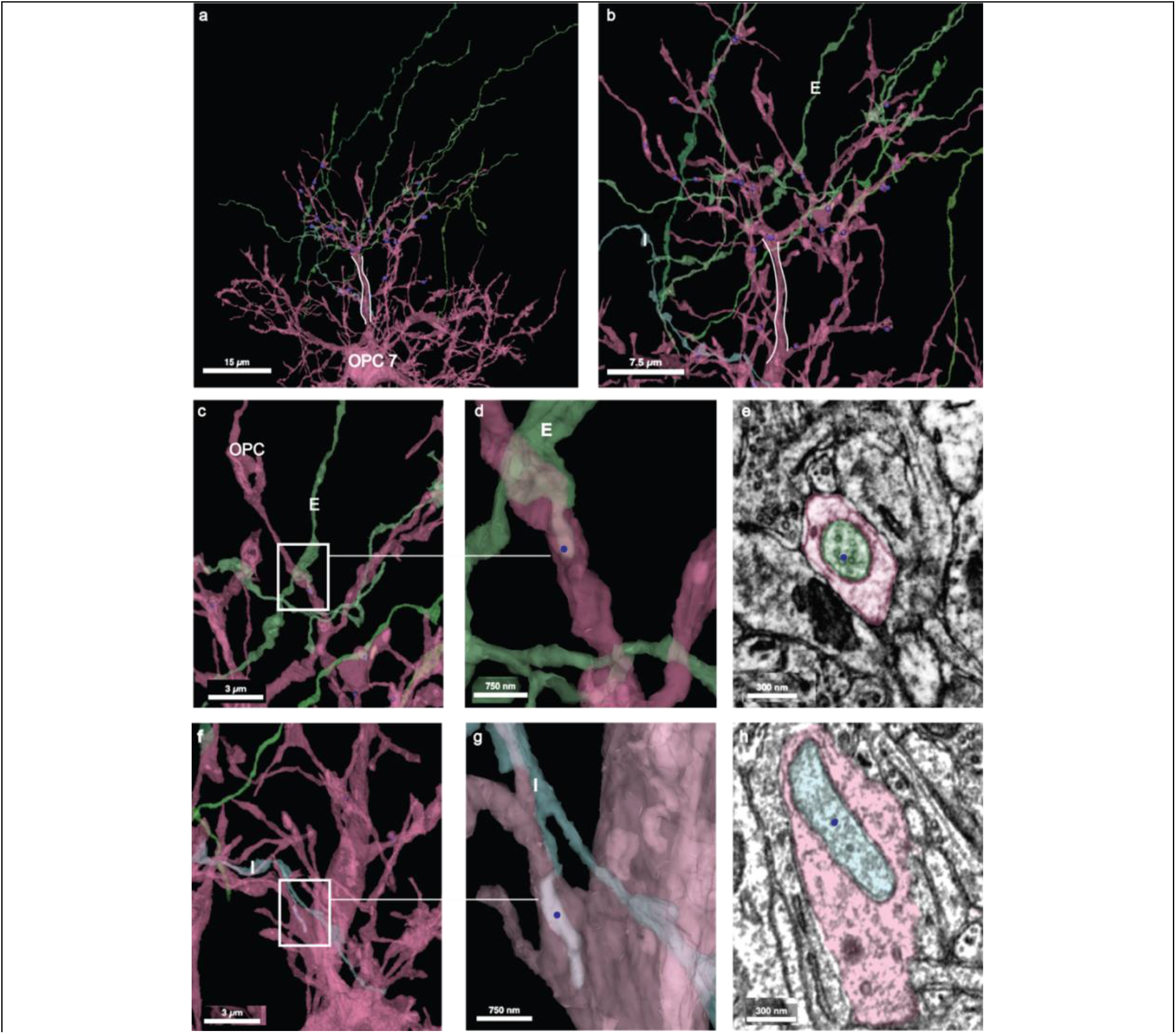
Axons engulfed within OPC7 main branch. **a)** One branch (while outline mark trunk) of OPC7(pink) was used to annotate axons engulfed within OPC cytoplasm. Scale bar, 15 μm. **b)**Higher magnification view of OPC branch trunk (white outline) and its processes(pink) and excitatory axons(E) in green. Scale bar, 7.5 μm. **c)** A portion of an excitatory axon(boxed area) is engulfed within the OPC cytoplasm(pink). Scale bar, 3 μm. **d,** Higher magnification view of boxed area shows engulfed tip (blue dot) of excitatory axon(E). Scale bar, 750 nm. **e,** Ultrathin section view of cross section of the green excitatory axon inside the OPC process(blue dot). Scale bar, 300 nm. **f,** A portion of an inhibitory (I) axon in blue (boxed area)is engulfed within OPC cytoplasm(pink). Scale bar, 3 μm. **g,** Higher magnification view of boxed area shows portion of the engulfed inhibitory axon branch (blue dot)within the OPC cytoplasm(pink). Scale bar, 750 nm. **h,** Ultrathin section view shows the inhibitory axon(blue dot) surrounded by OPC cytoplasm(pink). Scale bar, 300 nm.

